# Low soil moisture induces recruitment of Actinobacteria in the rhizosphere of a drought-sensitive and Rhizobiales in a drought-tolerant potato cultivar

**DOI:** 10.1101/2023.05.26.542284

**Authors:** Benoit R. Martins, Roberto Siani, Krzysztof Treder, Dorota Michałowska, Viviane Radl, Karin Pritsch, Michael Schloter

## Abstract

Growing evidence suggests that soil microbes can improve plant fitness under drought. However, in potato, the world’s most important non-cereal crop, the role of the rhizosphere microbiome under drought has been poorly studied. Using a cultivation independent metabarcoding approach, we examined the rhizosphere microbiome of two potato cultivars with different drought tolerance as a function of water regime (continuous versus reduced watering) and manipulation of soil microbial diversity (i.e., natural (NSM), vs. disturbed (DSM) soil microbiome). Water regime and soil pre-treatment showed a significant interaction with bacterial community composition of the drought-sensitive (HERBST) but not the drought-resistant cultivar (MONI). Depending on the cultivar, different taxa responded to reduced watering. Under NSM conditions, these were mostly rhizobiales order representative in MONI, and *Streptomyces*, *Glycomyces*, *Marmoricola*, *Aeromicrobium*, *Mycobacterium*, amongst Actinobacteriota, and the root endophytic fungus *Falciphora* in HERBST. Under DSM conditions and reduced watering, *Bradyrhizobium*, *Ammoniphilus*, *Symbiobacterium* and unclassified Hydrogenedensaceae responded in the rhizosphere of MONI compared to the continuous, while in HERBST, fewer taxa of Actinobacteriota and no fungi responded to reduced vs. continuous watering. Overall, our results indicate a strong cultivar specific relationship between potato and their associated rhizosphere microbiomes under reduced soil moisture.

## 1. Introduction

Drought is one of the major factor limiting crop productivity and ultimately food security (Fahad et al., 2017). Thus, drought-tolerance is one of the major aims in breeding programs for crops (Zhang et al., 2022). Potato is the most produced vegetable crop worldwide (FAOSTAT 2021), and among the hundreds of commercial cultivars, there is a considerable variation in drought tolerance (Sprenger et al., 2015, 2016; Stark et al., 2013). Since very short periods of water shortage can lead to substantial yield loss (Nasir and Toth, 2022), potato is generally considered drought-sensitive crop (Yuan et al., 2003). Drought susceptibility has been mainly attributed to its shallow root system (Monneveux et al., 2013; Obidiegwu et al., 2015) with weak soil penetration (Joshi et al., 2016; Stalham et al., 2007), but there are also complex environmental interactions (Spitters and Schapendonk, 1990). Thus, besides breeding efforts in potato (Tiwari et al., 2022), additional approaches particularly the use of plant growth promoting and fortifying microbes are considered an important component to improve plant health and performance (Berg, 2009; Ngumbi and Kloepper, 2016).

Positive effects of soil microbial diversity have been associated with soil multifunctionality (nutrient cycling, primary production, litter decomposition etc.) (Delgado-Baquerizo et al., 2016). Particularly, microbes inhabiting the plant rhizosphere and their complex interactions with the host plant significantly affect plant morphology, physiology, plant growth, development, and health (Philippot et al., 2013). Therefore, plant-microbe interactions are thought to play a critical role in the fast adaptation of plants to environmental stress conditions (Pascale et al., 2020).

Considered as the second plant genome (Berendsen et al., 2012; Li et al., 2021; Yang et al., 2022), the rhizosphere microbiome contributes to a broad spectrum of functions including plant nutrition, defence against phytopathogens, and adaptation to abiotic and biotic stresses (Garcia and Kao-Kniffin, 2018; Vandenkoornhuyse et al., 2015). Especially, as a response to drought, plants significantly shift their rhizosphere microbiome by recruiting drought tolerant taxa (Naylor et al., 2017; Santos-Medellín et al., 2017; Xu et al., 2018). Reversely, rhizosphere microbial populations are influenced by a complex combination of factors. Among these, soil acting as microbial seed bank plays the most significant role in determining the composition of rhizosphere microbial communities as shown for populus (Veach et al., 2019), cotton (Yang et al., 2022) and soybean (Liu et al., 2019), as well as for potato with cultivars grown in two different soils recruited different rhizosphere microbiomes (Inceoǧlu et al., 2012).

Moreover, rhizosphere microbial composition is shaped by host species, as shown by the recruitment of different communities in the same soil by different plant species (Berg et al., 2006; Garbeva et al., 2008). Additionally, cultivars within a single plant species were shown to acquire different rhizosphere communities (Fan et al., 2023; Q. Liu, Xie, et al., 2021; Micallef et al., 2009). Among the possible structuring factors, in addition to root morphology, root exudates are considered the main drivers in shaping microbiome in the rhizosphere (Bulgarelli et al., 2013). The exudation pattern (quantity and quality), which depends on the genotypes (Badri and Vivanco, 2009; Gargallo-Garriga et al., 2018), have been showed to vary amongst potato cultivars (Gschwendtner et al., 2011).

The rhizosphere microbiome can also be affected by environmental factors including drought. Drought-induced transformations in the microbiome have been attributed to changes in plant physiology and biochemistry, which compromise carbon compound efflux and root exudate profiles (Chen et al., 2022; Gargallo-Garriga et al., 2018; Naylor and Coleman-Derr, 2018). Nevertheless, different microbial groups in the rhizospheres of various plants exhibited different tolerances towards reduction in soil moisture. A study of drought effects on root associated microbiomes of 18 grass species showed that drought consistently enriched Actinobacteria lineages among hosts, but depleted Acidobacteria and Betaproteobacteria (Naylor et al., 2017). Furthermore, the enrichment was more pronounced as the plant-microbe interactions increased (endosphere > rhizosphere > surrounding soil) (Naylor et al., 2017), implying that Actinobacteria play an important role in drought resistance of plants (Naylor et al., 2017). Actinobacterial enrichment was also observed in sorghum (Xu et al., 2018), tall wheatgrass (Naylor et al., 2023) and rice (Santos-Medellín et al., 2017, 2021), and root colonization of the Actinobacterial genus *Streptomyces* was consistent with increases in root growth under drought (Xu et al., 2018). This provides evidence that specific microbial taxa can support plant growth under drought. However, diverse mechanisms behind microbial-mediated drought resistance in plants have been previously reported, notably those modulating phytohormones levels in stressed plants (Bhattacharyya et al., 2021). This includes mechanisms reducing levels of the plant stress hormone ethylene by amino-1-cyclopropane carboxylate (ACC) deaminase-producing bacteria (Glick, 2014), particularly in drought-enriched Actinobacteria (Gebauer et al., 2022). The microbial production of auxins has also been suggested to improve root traits related to drought tolerance such as root length, root tips and surface area (Jochum et al., 2019).

There is no clear pattern on how drought affects fungal communities in the soil and in the rhizosphere. While many studies showed no or minor effects of drought on fungal community composition (Barnard et al., 2013; Furze et al., 2017; Naylor et al., 2017; Ochoa-Hueso et al., 2018), other studies found significant changes under drought (Bazany et al., 2022; Carbone et al., 2021; Santos-Medellín et al., 2017).

In potato, little is known on how drought affects the interactions between cultivars of different drought tolerance (Boguszewska-Mańkowska et al., 2020; Sprenger et al., 2015; Stark et al., 2013), and their rhizosphere microbiomes. The aim of this study was to investigate the potential role of the rhizosphere microbiome in sustaining potato growth under reduced soil moisture. We performed a greenhouse experiment with two potato cultivars of different drought tolerance and cultivated the plants in soil with a natural microbiome, and in soil with an artificially reduced microbiome. We compared the rhizosphere microbiome (bacteria, fungi) of each cultivar grown under optimal and reduced watering. Given that plants interact with a plethora of microorganisms at the roots to form unique rhizosphere microbial associations that respond to environmental conditions (Pascale et al., 2020), we hypothesized that when drought tolerance is mediated in potato by the recruitment of beneficial microbes, the rhizosphere microbiome would be affected in a cultivar-specific manner under reduced soil moisture, with the more drought resistant cultivar exhibiting a more drought adaptive microbiome than the drought susceptible cultivar (**H1**).

For the soil with an artificially reduced microbiome, we hypothesized that when the rhizosphere microbiome plays a dominant role in drought resistance of the two potato cultivars, then a reduction of the soil microbiome would result in lower drought tolerance under reduced watering in both cultivars (**H2**).

## 2. Material and Methods

### 2.1 Soil sampling and analysis of physico-chemical properties

In spring 2020, top of luvisol (0-20 cm) characterised as a silty loam was obtained from the site Gut-Roggenstein experimental station (Olching, Bavaria, Germany), Technical University of Munich. The soil contained 1.27% total carbon and 0.1% total nitrogen resulting in a C:N ratio of 12.7. Previously, the site was used consecutively for the cultivation of different crops such as summer barley in 2015, sugar beet in 2016, rapeseed in 2017, wheat in 2018, winter barley in 2019 and sugar beet in 2020. The soil was sieved with a 2-mm diameter mesh and divided into two portions. One was autoclaved for 20 min at 121 °C (to reduce soil microbial biomass and diversity) and the other was left in its native state. Before potting and planting, both soils were stored at 8°C for 7 days.

Bare NSM and DSM soils (non-planted soils) were characterised for microbial biomass carbon (C_mic_) and nitrogen (N_mic_), using chloroform fumigation-based extraction method (Joergensen, 1996; Vance et al., 1987), which were calculated as the difference between total dissolved organic carbon (DOC) and nitrogen (DON) in fumigated and non-fumigated samples, with an extraction efficiency coefficient (k_ec_) value of 0.45 (Vance et al., 1987) for carbon and k_en_ value of 0.54 (Brookes et al., 1985) for nitrogen. Inorganic nitrogen (NH_4_-N and NO_3_-N), pH, soil texture (clay, silt and sand contents), magnesium (Mg), phosphorus (P_2_O_5_) and potassium (K_2_O) as well as the maximum water holding capacity (mWHC) were also determined using standard protocols. Details on soil characteristics and protocols used are summarised in the supplementary document.

### 2.2 Experimental design

Two potato (*Solanum tuberosum*) cultivars were used in this experiment (**Figure 1**), MONI referenced as drought-resistant and HERBSTRFREUDE (HERBST used throughout the text) considered as drought-sensitive (Personal communication of Krzysztof Treder). Additional information about the two cultivars is available on the European Cultivated Potato Database (https://www.europotato.org/). Plants of the two cultivars were vegetatively propagated as tissue cultures at the Institute of Plant Breeding and Acclimation in Bonin (Bonin, Poland). Plant tissues were grown in test tubes under *in vitro* conditions using Murashige and Skoog nutrient medium for approximately 8 weeks. Certified healthy plants (pathogen-free) were delivered to Helmholtz Munich (Munich, Bavaria, Germany). Agar plugs attached to the roots of the plantlets were gently removed with tweezers and tap water. Plantlets were transferred to 0.3 L (7×7×8 cm) pots filled with NSM or DSM soils and allowed to acclimate for 2 weeks. During the acclimation period, the early-stage plants were watered thrice per week.

**Figure 1.**
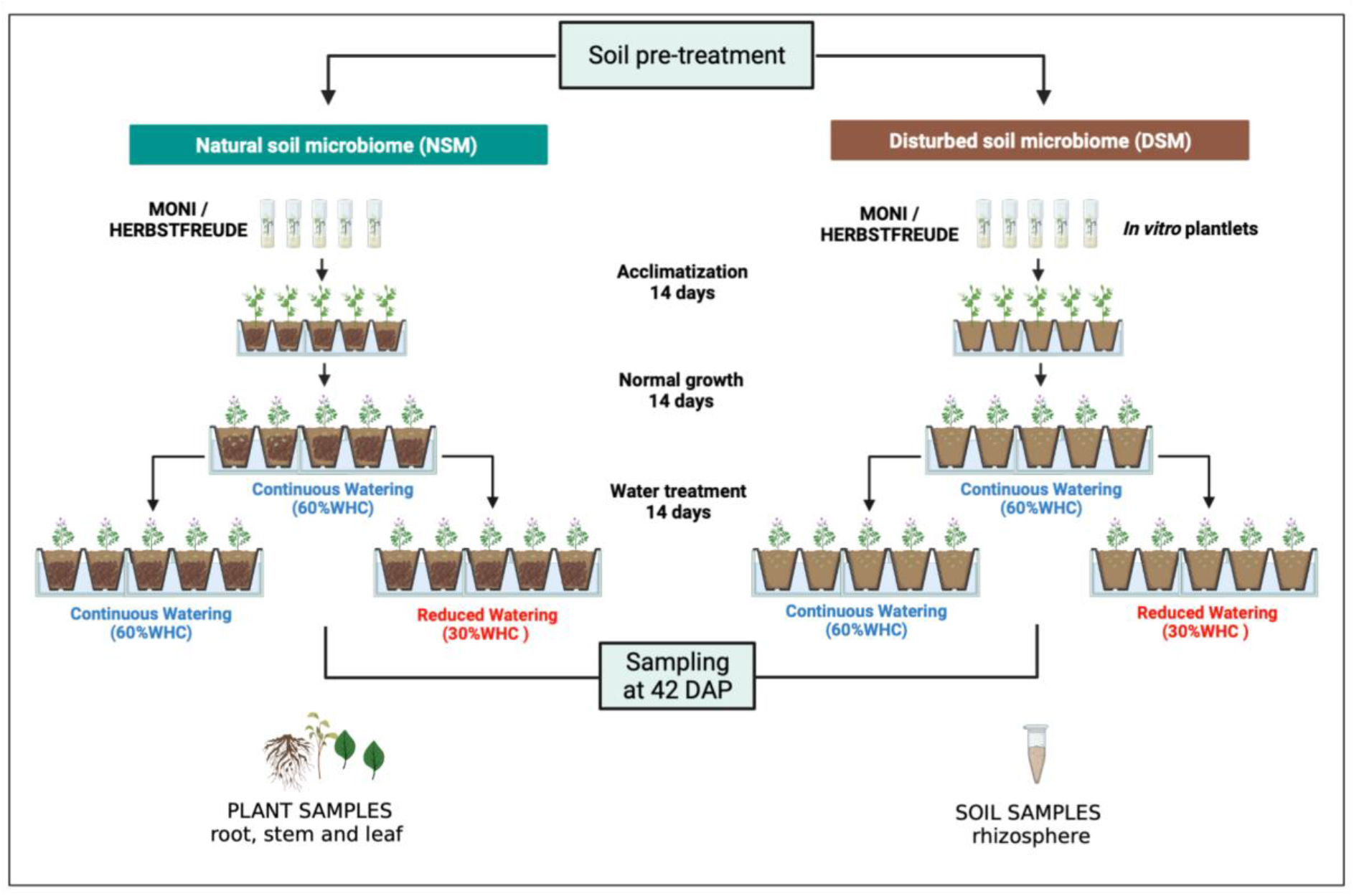
Experimental design of the greenhouse experiment using two *in-vitro* propagated potato cultivars (MONI, HERBSTFREUDE) acclimated on soil with natural (NSM) and disturbed soil microbiome (DSM) for 14 days. After transfer to 1.5 L pots containing the corresponding soils, plants were grown for another 14 days under control watering (60% water holding capacity (WHC)). After 28 days, half of the plants were kept under 60% WHC, the other half were kept under reduced watering (30% WHC). Upon harvest at day 42, plants samples (root, stem, leaves) were taken to assess growth traits, and rhizosphere soil samples were collected for bacterial and fungal community analyses. Created with BioRender.com

The acclimated plants were afterwards transferred to 1.5 L (11×11×12 cm) pots, where they were grown for 14 days under optimal greenhouse conditions (RH: 65%; day/night temperature: 22°C/18°C day/night natural photoperiod: 16/8) and were maintained with continuous watering (CW = 60% of maximum soil water holding capacity). Subsequently, plants grown either in NSM or DSM soils were divided in two groups, one was continuously kept under continuous watering whereas the second was maintained with reduced watering (RW = 30% maximum soil WHC). The entire trial was conducted without applying fertilisers to not influence the soil microbiome. Three unplanted pots each filled with NSM and DSM soils served as controls for water content adjustments throughout the experiment. 14 days after plants were imposed to the two water regimes, the experiment was completed. Plant growth parameters including stem height, leaf dry mass, root length and fresh weight were measured. The soil adhering to the plant roots, defined as the rhizosphere, was collected per plant. This three-factorial experiment (2 soil pre-treatments x 2 water regimes x 2 cultivars x 5 replicates) resulted in 8 experimental treatments and 40 rhizosphere samples. After autoclaving and prior to planting, NSM and DSM soils were sampled for assessment of the initial soil microbiome (T0). Bare soils and rhizosphere samples were stored at -80°C until DNA extraction and library preparation for amplicon sequencing.

### 2.3 DNA Extraction and library preparation for amplicon sequencing

DNA was extracted from 450 mg of soil samples taken before planting (T0) and from the rhizosphere taken at day 42 (harvest), according to Griffiths et al., (2000). Empty extraction tubes were used as negative controls to check for contamination during the process. The concentration of total DNA extracts was quantified in duplicate using SpectraMax Gemini EM Microplate Spectrofluorometer (Molecular Devices, CA, USA) and Quant-iT PicoGreen dsDNA assay kit (Thermo Fischer Scientific, Waltham, USA) according to the manufacturer’s instructions. All samples were stored at -20°C until further analysis.

The ITSmix3/ITSmix4 primer pair (Tedersoo et al., 2014) was used to amplify the ITS2 region of the fungal nuclear rDNA. Unicate PCR was performed with an initial denaturation phase at 95°C for 10 min and 30 cycles of 30s denaturation at 95°C, 30s annealing at 55°C and 1 min extension at 72°C and a final extension of 10min at 72°C. Amplification of the V4 region of the bacterial 16S rRNA gene required the use of universal primer pair 515F/806R (Apprill et al., 2015; Parada et al., 2016). Unicate PCR was performed under the following conditions: an initial denaturation phase at 98°C for 1 min and 30 cycles of 10s denaturation at 98°C, 30s annealing at 55°C and 30s extension at 72°C and a final extension for 5 min at 72°C. ITS and 16S PCR were carried out in the following reaction mixture: 1x NEBNext High Fidelity Master Mix (New England Biolabs, Ipswich, USA), 5 pmol of each primer, 3% bovine serum albumin (BSA), 5 ng of template DNA, and DEPC water to a final volume of 25 μL.

PCR products were verified in 1% agarose gels, followed by MagSi NGSprep Plus bead purification (Steinbrenner, Wiesenbach, Germany). The quality and quantity of purified amplicons and the presence of primer dimers were checked with DNF-473 Standard Sensitivity NGS Fragment Kit (1-6000 bp) on a fragment analyser (Agilent Technology, Santa Clara, California, USA). DNA concentrations were adjusted to 2ng/μL. 8-cycle indexing PCR was performed in a reaction mix (25 μL) using 2.5 μL of each indexing primer (Nextera® XT Index Kit v2; Illumina, San Diego, California, United States), 12.5 μL NEBNext High-Fidelity 2× PCR Master Mix, 1.5 μL DEPC-treated water and 10 ng of purified amplicon. Indexed amplicons were subjected to a second round of purification with quality and quantity as described previously. Prior to sequencing, samples were normalised to 4nM and equimolarly pooled into a single Eppendorf tube. Paired-end sequencing was carried out using the MiSeq ® Reagent kit v3 (600 cycles) (Illumina Inc., San Diego, California, USA).

### 2.4 Pre-processing of sequencing data

Raw sequencing data were analysed using the Galaxy web platform (www.usegalaxy.org; Afgan et al., 2016). After the raw data were imported into the platform, forward and reverse FASTQ files were used to build their respective dataset lists. Forward and reverse dataset lists were trimmed with a minimum read length of 50 using the Cutadapt function (Martin, 2011). Quality control for the forward and reverse reads was performed via FastQC (Andrews, 2010). Subsequently, data analysis was performed using the DADA2 pipeline (Galaxy Version 1.20) (Callahan et al., 2016). The following trimming and filtering parameters were considered for 16S rRNA analysis: 20 bp were removed n-terminally and reads were truncated at position 230 (forward) and 180 (reverse) with expected error of 3 and 4, respectively. For the ITS2 analysis, forward reads were trimmed to 20-220 bp, reverse reads to 20-160 bp and same number of expected errors was used. The remaining reads were merged and further inferred into unique amplicon sequence variants (ASVs).

ASVs are biological sequences discriminated from errors, allowing the detection of single-nucleotide differences over the sequenced genes. For bacterial taxonomic assignment, ASVs were trained against SILVA database v138.1 with 0.99 confidence threshold. Fungal ASVs were taxonomically assigned using UNITE fungi database v9.0 released for QIIME with 0.99 confidence threshold (Abarenkov et al., 2022). Amplicon sequences from bacteria and fungi were aligned and phylogenetic trees were constructed. The R language and environment v4.2.1 were used for downstream analysis. Using Bioconductor decontam package v1.13.0 (Davis et al., 2018), contaminant sequences were filtered leveraging the negative controls, along with ASVs assigned to chloroplast and mitochondria. A phyloseq object was created for each of bacterial and fungal datasets using the Phyloseq package v1.42.0. Singletons (ASVs represented by only one read across all samples) were removed. Furthermore, only ASVs consistently found in 80% of the biological replicates (4 out of 5) in each sample collection were kept for downstream analysis. A summary report on the phyloseq objects is available in the supplementary table S3. For normalization, Total-Sum Scaling (TSS) which transforms the abundance table into relative abundance table, i.e., scale by each sample’s library size was applied.

### 2.5 Statistical analyses

Bare NSM and DSM soils were characterised regarding physico-chemical properties and statistical differences were calculated with a student t-test. The normality of the distribution in each group as well as equality of variance were checked using Shapiro-Wilk and F-test, respectively.

Statistical differences between plant growth parameters in different soil pre-treatments were calculated using the non-parametric sum rank Wilcoxon test. Throughout the text, the significance level for all statistical tests was set at 0.05.

For α-diversity estimation of bacterial and fungal communities in the plant rhizosphere, observed species richness and Shannon index were calculated across sample groups and visualized using the Microbiome v1.20.0 and ggplot2 v3.4.0, respectively. Wilcoxon rank sum test was conducted to assess the effect of soil pre-treatment and water regimes on the α-diversity. To partition the source of variance, the relative contribution of soil pre-treatment, and water regimes was assessed using a permutational multivariate analysis of variance (PERMANOVA) based on UniFrac dissimilarity matrices as implemented in the adonis2 function (R package vegan v2.6-4).

The analysis of differentially abundant taxa across sample groups was conducted with LEfSe as implemented in the diff_analysis function (Bioconductor R package MicrobiotaProcess, v1.10.2) under one-against-all mode (i.e., one taxon is significantly different only when it is significantly different against all remaining treatments). In brief, a Kruskal-Wallis test, followed by a Wilcoxon signed-rank test were used to isolate differentially abundant features. For overall abundance comparison between continuous and reduced watering across all microbial taxa, logarithmic LDA score threshold set to 2.5 was calculated and any taxa with α less than 0.05 were defined to be significantly different between water regimes. To provide a comprehensive comparison of microbial abundance across sample groups, the LEfSe analysis between continuous and reduced watering samples was performed at genus level.

Comparative Venn diagram analysis was performed to identify overlapping ASVs between the two cultivars amongst soil pre-treatment and water regimes. The shared microbiomes under the different treatments were defined using ASVs that were found in 80% of the replicates in each sample group with a relative abundance threshold of 0.001. The analysis was computed with the function amp_venn of the package ampvis2 (v2.7.34). The function amp_heatmap from the same package was used to visualise the shared microbiome composition and their relative abundances.

## 3. Results

### 3.1 Plant growth parameters

Plant growth parameters revealed that soil pre-treatment and water regimes had no significant effect on root length of the cultivar MONI, whereas HERBST exhibited a significant reduction of its root length when grown in DSM soil under reduced watering (**Figure 2A**). Aboveground, both treatments did not affect stem height in either cultivar (**Figure 2B**). Surprisingly, MONI cultivated with reduced watering in DSM soil showed a significantly higher leaf dry weight compared to NSM soil (**Figure 2C**) but no change in leaf dry weight of HERBST was observed.

**Figure 2.**
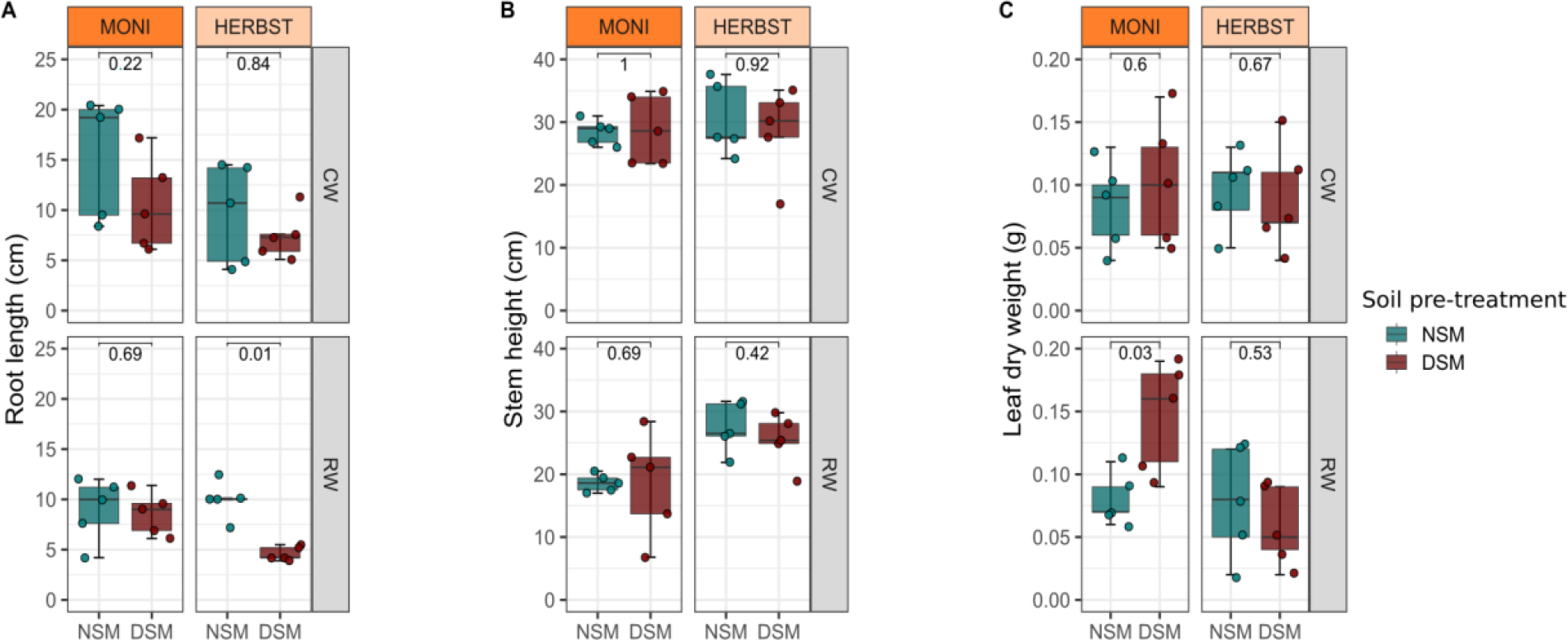
Comparison of plant parameters between the cultivars MONI and HERBST grown in soil with natural microbiome (NSM, green bars) or autoclaved soil with disturbed microbiome (DSM, brown bars) under continuous (CW) and reduced (RW) watering regimes A) roo t length B) stem height and C) leaf dry weight Horizontal bars within boxes are the median. The tops and bottoms of the boxes represent 75th and 25th quartiles, respectively. The two vertical lines outside the boxes represent the whiskers. The colored dots stand for the individual observations. A non-parametric Wilcoxon sum-rank test (p < 0.05, n = 5) was applied to calculate significant differences across sample groups and numbers above the boxes indicate the statistical p-values.

### 3.2 Effect of soil pre-treatment and water regimes on diversity and composition of rhizosphere microbiomes

As expected, pre-treatment of soil resulted in lower bacterial α-diversity of the rhizosphere microbiome in DSM soil, and in rhizosphere samples from both cultivars, MONI (Observed: Wilcoxon rank sum test, p = 0.00013) and HERBST (Observed: Wilcoxon rank sum test, p = 2.2e-05) (**Table 1A**). In comparison to the continuous, reduced watering had no significant effect on bacterial α-diversity of MONI, but consistently resulted in higher diversity in the cultivar HERBST (**Figure 3A**).

**Figure 3.**
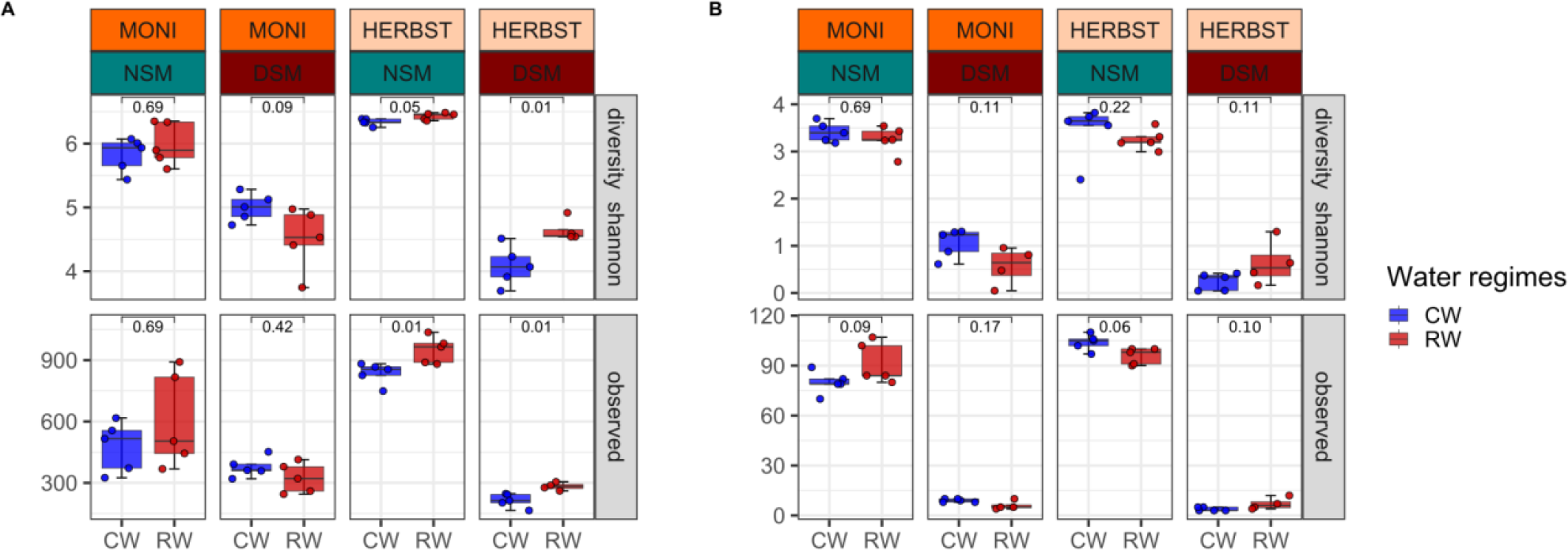
Comparison of α-diversity based on Shannon index and observed species of A) bacterial and B) fungal communities in rhizosphere soil samples of the potato cultivars MONI and HERBST grown in either native soil with natural microbiome (NSM) or autoclaved soil with disturbed microbiome (DSM), and either continuous watering (blue) or reduced watering (red). Boxplots display the medians, tops and bottoms of the boxes represent 75th and 25th quartiles, and whiskers outside this range; dots illustrate the individual observations in each sample group. A non-parametric Wilcoxon sum-rank test (p < 0.05, n = 5) was applied to calculate significant differences across sample groups and numbers above the boxes indicate the corresponding p-values.

**Table 1.** Effect of soil pre-treatment (natural (NSM) vs. disturbed (DSM) soil microbiome) on α-diversity metrics (Shannon index, observed species) of A) bacterial and B) fungal communities in the rhizosphere of two potato cultivars (MONI, HERBST). A non-parametric Wilcoxon sum-rank test (p < 0.05, n = 10) was applied to calculate significant differences across sample groups.

| Cultivars | Metrics | group1 | group2 | p | p.adj | p.format | p.signif | method |
| --- | --- | --- | --- | --- | --- | --- | --- | --- |
| MONI | Shannon | NSM | DSM | 1.1E-05 | 1.1E-05 | 1.10E-05 | **** | wilcoxon |
| MONI | observed | NSM | DSM | 0.00684 | 0.0068 | 0.0068 | ** | wilcoxon |
| HERBST | Shannon | NSM | DSM | 0.0000217 | 0.000022 | 2.20E-05 | **** | wilcoxon |
| HERBST | observed | NSM | DSM | 0.0000217 | 0.000022 | 2.20E-05 | **** | wilcoxon |

**B**
| Cultivars | Metrics | group1 | group2 | p | p.adj | p.format | p.signif | method |
| --- | --- | --- | --- | --- | --- | --- | --- | --- |
| MONI | Shannon | NSM | DSM | 0.0000217 | 0.000022 | 2.20E-05 | **** | wilcoxon |
| MONI | observed | NSM | DSM | 0.000266 | 0.00027 | 0.00027 | *** | wilcoxon |
| HERBST | Shannon | NSM | DSM | 0.0000217 | 0.000022 | 2.20E-05 | **** | wilcoxon |
| HERBST | observed | NSM | DSM | 0.000264 | 0.00026 | 0.00026 | *** | wilcoxon |

Soil pre-treatment significantly affected fungal α-diversity, being lower in DSM soil for the two cultivars, MONI (Observed: Wilcoxon rank sum test, p = 0.00027) and HERBST (Observed: Wilcoxon rank sum test, p = 0.00026) (**Table 1B**) whereas the effect of the two water regimes was insignificant (**Figure 3B**).

PERMANOVA test revealed that soil pre-treatment significantly altered bacterial composition in the rhizosphere of both cultivars, MONI (weighted UniFrac, R^2^ = 0.895, p = 0.001), and HERBST (weighted UniFrac: R^2^ = 0.866, p = 0.001). In contrast, water regimes did not affect the bacterial composition in MONI (weighted UniFrac: R^2^ = 0.011, p = 0.17) however, we found a significant interaction of water regimes with soil pre-treatment in HERBST (weighted UniFrac: R^2^ = 0.021, p = 0.053). Fungal community was also driven by soil pre-treatment in both cultivars, MONI (weighted UniFrac, F = 9.7419, p = 0.001) and HERBST (weighted UniFrac, R^2^ = 0.647, p = 0.001) but not influenced by water regimes.

### 3.3 Microbial responders

Linear Discriminant Analysis Effect Size (LEfSe) revealed that rhizosphere bacteria and fungi identified as responders significantly differed across water regimes, cultivars, and pre-treatment of the soil (**Figure 4**). For MONI, Proteobacteria (unclassified Rhizobiales and Rhodanobacteraceae), Verrucomicrobia (Ellin 517) and Actinobacteriota (unclassified Microtrichales) were identified as potential responders in reduced watering samples from NSM soil whereas those were Proteobacteria (*Bradyrhizobium*), Bacillota (*Ammoniphilus*)*, Firmicutes* (*Symbiobacterium*) and Hydrogenedentes (unclassified Hydrogenedensaceae) from DSM soil (**Figure 4A**). For HERBST grown in NSM soil, responders to the reduced watering included mainly Actinobacteriota lineages (*Streptomyces, Marmoricola, Aeromicrobium, Glycomyces, Mycobacterium,* unclassified Acidimicrobiia *and 0319-7L14*) and a Myxococcota (unclassified Sandaracinaceae). In DSM soil, although potential responders of reduced watering included Actinobacteriota (*Marmoricola*, *Nocardioides*, Acidimicrobiia unclassified), de novo taxonomic groups such as Proteobacteria (*Novosphingobium, Sphingobium, Hirshia, Caenimonas*, unclassified Devsociaceae and Comamonadaceae), Myxococcota (*Sandaracinus*) and Acidobacteriota (unclassified Vicinamibacterales and Vicinamibacteraceae) were also observed (**Figure 4A**). With respect to fungal communities, potential responding taxa associated with the reduced watering in NSM soil consisted of Ascomycota (*Magnaporthiopsis,* and unclassified Helotiales) in MONI and the genera *Falciphora* and *Neocosmospora* in HERBST (**Figure 4B**). For both cultivars, no biomarkers of water regimes were identified in DSM soil.

**Figure 4.**
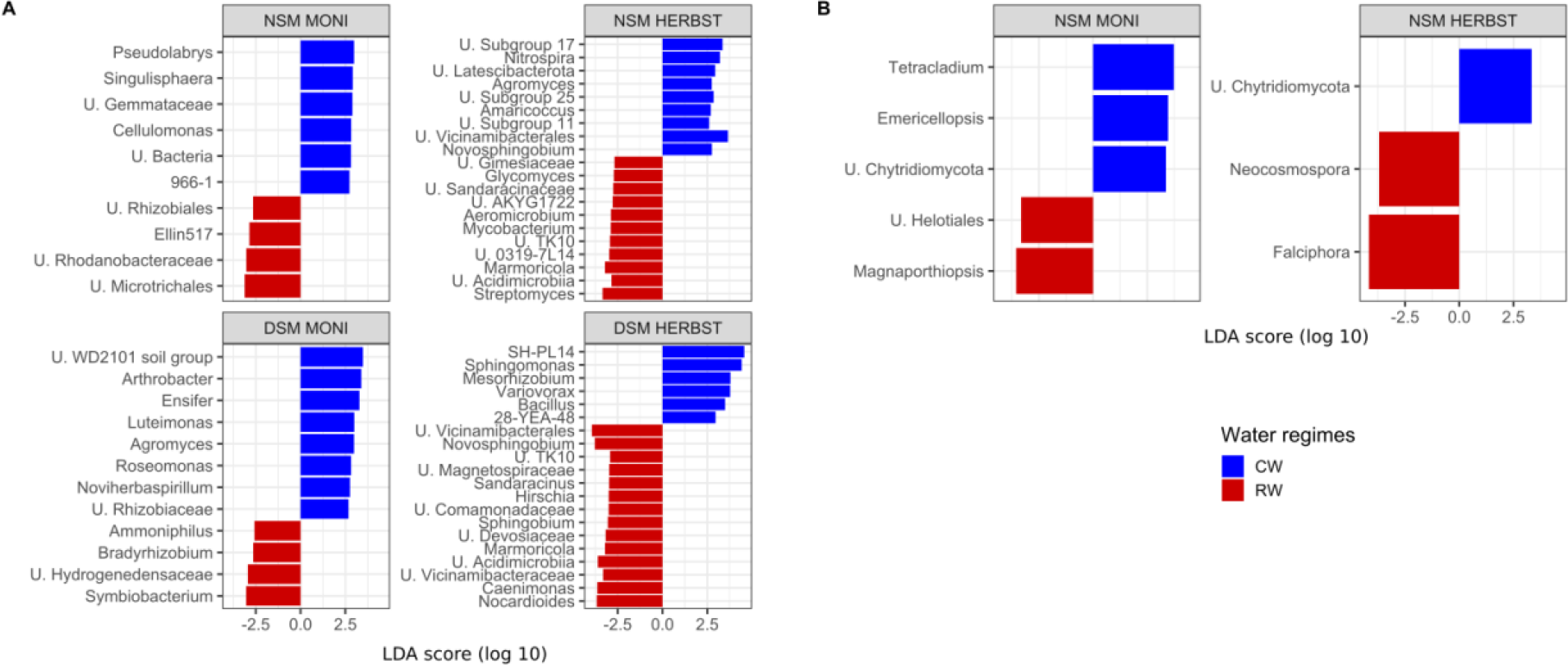
Linear Discriminant Analysis (LDA) combined with Effect Size (LEfSe) plot of A) bacterial and B) fungal genera identified as potential biomarkers for different watering regimes in the rhizosphere of the potato cultivars MONI and HERBST grown in native so il with natural microbiome (NSM) and autoclaved soil with disturbed microbiome (DSM), respectively. p-values < 0.05 were considered significant for factorial Kruskal–Wallis and pairwise Wilcoxon tests. Colors refer to the watering regimes (blue continuous watering, red reduced watering). Only the 20 most differentially abundant features meeting an LDA significant threshold ≥ 2.5 are shown.

### 3.4 Shared microbiomes across soil pre-treatment and water regimes

In NSM soil, both cultivars shared the highest amount of bacterial ASVs under reduced watering, i.e., 342 ASVs representing 67.8% of the total read count (**Figure 5B**), compared to 264 ASVs (62.3%) under continuous watering (**Figure 5A**). Phylum-based analysis revealed a similar composition of the microbiomes shared by the two cultivars under continuous watering and reduced watering, but slight changes occurred in the relative abundances (**Figure 6A-B**). Actinobacteriota were by far the most abundant phylum followed by Proteobacteria, Acidobacteriota and Chloroflexi. Compared to continuous watering, a slight increase and decrease in the proportions of Actinobacteriota and Acidobacteriota, respectively, was observed under reduced watering, whereas Proteobacteria and Chloroflexi remained relatively stable (**Figure 6A-B**). At the genus level, *Gaiella*, *Arthrobacter, Nocardioides* and unclassified Gaiellales, MB-A2-108, 67-14, were the most represented Actinobacteriota; *Hyphomicrobium* and Ellin6067 in Proteobacteria; unclassified Vicinamibacterales and Vicinamibacteraceae in Acidobacteriota and unclassified KD4-96, Gitt-GS-136, JG30-KF-CM45 in Chloroflexi (**Supplementary Figure S3A-B**). In DSM soil, MONI and HERBST also shared a microbiome under continuous watering (109 ASVs for 63% total read count) and reduced watering (115 ASVs for 65.3 % total read count) (**Figure 5C-D**).

**Figure 5.**
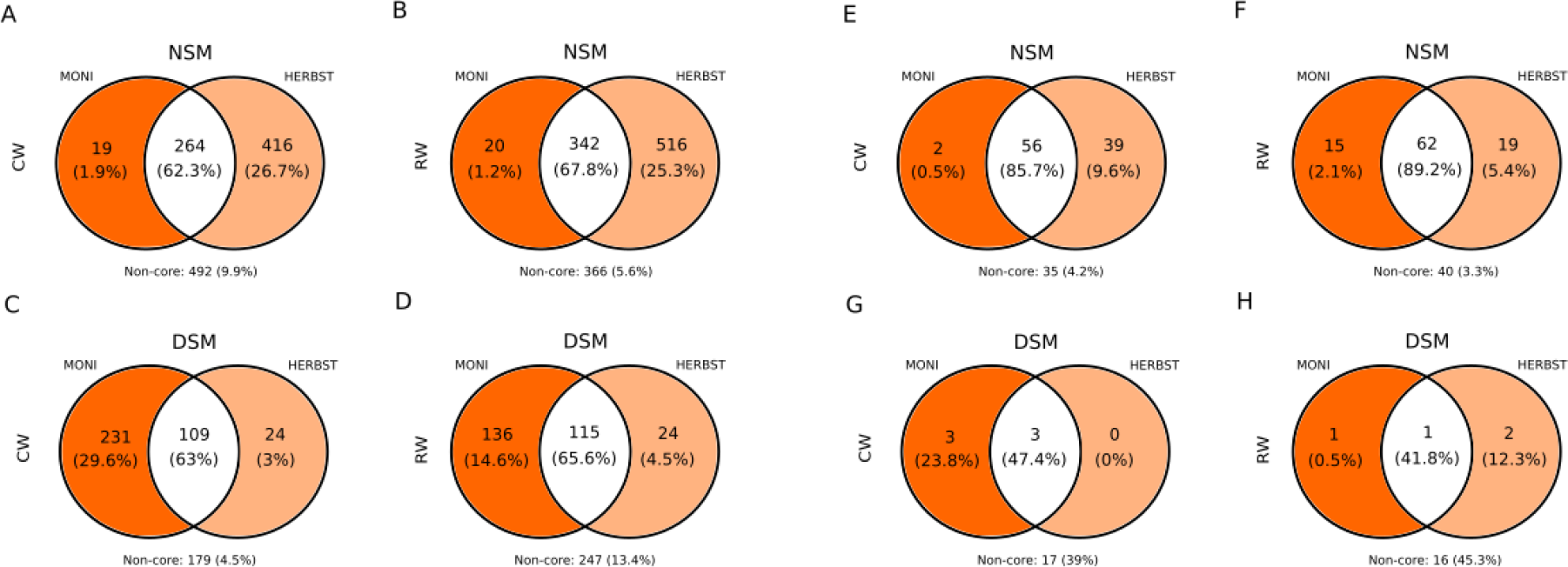
Venn diagram showing the number of unique (colored ellipses) and shared (white ellipses) bacterial and fungal ASVs to the cultivars MONI and HERBSTFREUDE grown in native soil with natural microbiome (NSM) and autoclaved soil with disturbed microbiome (DSM), under continuous (CW; A, C, E, G) and reduced (RW; B, D, F, H) watering. Respective read percentages are indicated in the brackets. Only ASVs found in the 80% of each sample collection with relative abundance of 0.001 were considered for analysis. Non-core represents the part of ASVs and percentage of reads not included in the frequency and abundance cut-offs.

**Figure 6.**
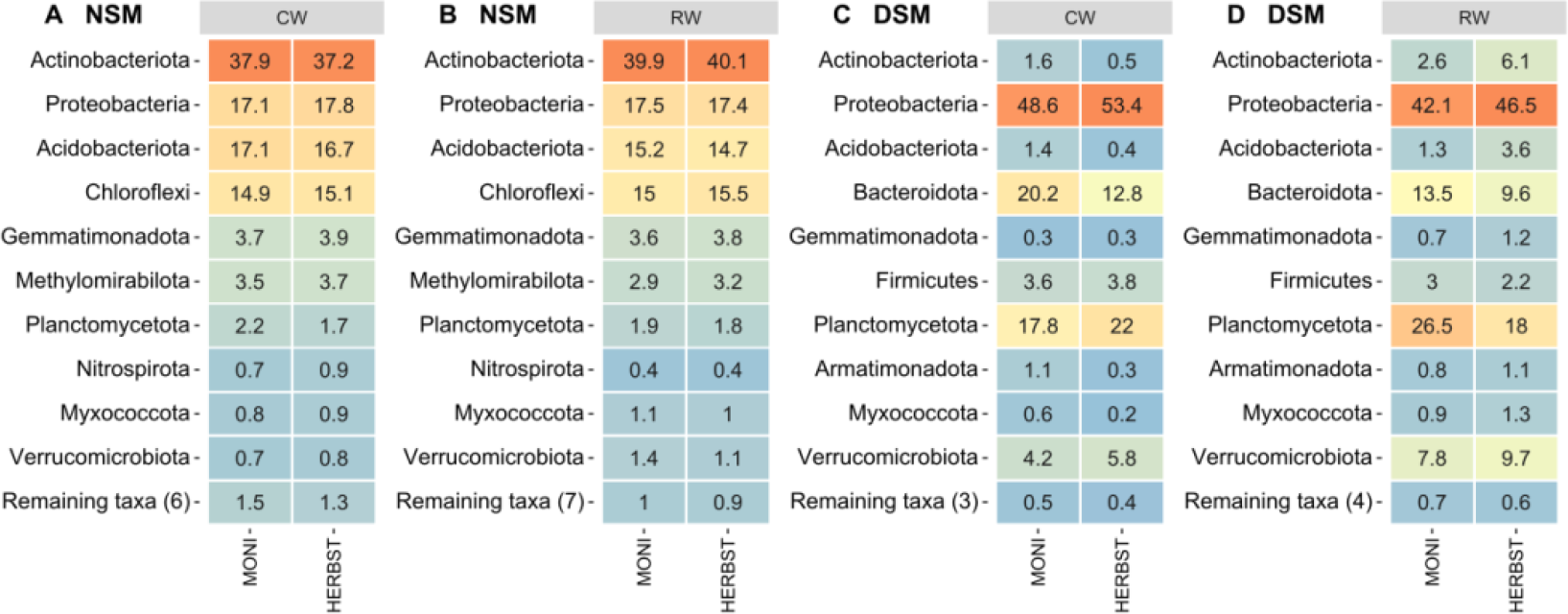
Bacterial shared microbiome composition. Heatmap displaying the top 10 phyla in the shared microbiome of cultivars MONI and HERBSTFREUDE grown in native soil with natural microbiome (NSM) and autoclaved soil with disturbed microbiome (DSM), under continuous (CW; A, C) and reduced (RW; B, D) watering. Numbers in the heatmap indicate the relative abundance of each taxon across sample collections.

Soil pre-treatment overall changed the composition of the shared microbiome at the phylum level in DSM soil by increasing in the relative abundances of Proteobacteria, Planctomycetes, Bacteroidota, Firmicutes and Verrucomicrobia at the expense of Actinobacteriota, Acidobacteriota, Chloroflexi and Gemmatimonadota (**Figure 6C-D**). For HERBST cultivated in DSM soil, Actinobacteriota increased under reduced watering, but their proportion remained marginal compared to the corresponding treatment in NSM soil. At the genus level, SH-PL14 was differentially abundant, especially enriched in MONI and depleted in HERBST under reduced watering treatment. *Sphingomonas* also decreased in HERBST under reduced watering but not in MONI (**Supplementary Figure S3C-D**). Comparing the two cultivars, the number of core ASVs was relatively stable in MONI regardless of the soil pre-treatment (**Table 2**), while in HERBST, the number of core ASVs was considerably lower in DSM soil compared to NSM soil (**Table 2**).

**Table 2.**
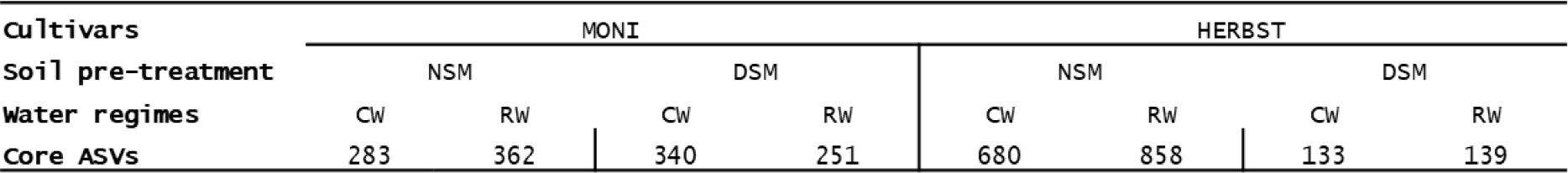
Summary of bacterial core ASVs in the rhizosphere of the cultivar MONI and HERBST across soil pre-treatment and water regimes.

The two cultivars grown in NSM soil had overlapping fungal ASVs with slightly more under reduced watering (62 ASVs for 89.2 % total read count) than continuous watering (56 ASVs for 85.7 % total read count) (**Figure 5E-F**). Shared ASVs were for the most assigned to the phylum Ascomycota (**Supplementary Figure S4**). At the genus level, *Podospora*, *Trichoderma* and *Falciphora* were among the top 10 taxa under reduced watering, but not continuous watering (**Figure 7A-B**). Interestingly, *Podospora* and *Falciphora* were not detected under continuous watering (**Supplementary Figure S4E**). *Phialophora* increased in proportion under reduced watering compared to continuous watering in both cultivars, whereas, *Gibellulopsis* remained stable in HERBST, but decreased in MONI under reduced watering relative to continuous watering (**Figure 7A-B**). In DSM soil, shared fungal ASVs were strongly reduced irrespective of water regimes (**Figure 5G-H**). Under continuous watering, 3 ASVs were shared, belonging to the genera *Acremonium* and *Exophiala*; under reduced watering, it was only 1 ASV, namely *Acremonium* (*Figure 7C-D*), all belonging to the phylum Ascomycota.

**Figure 7.**
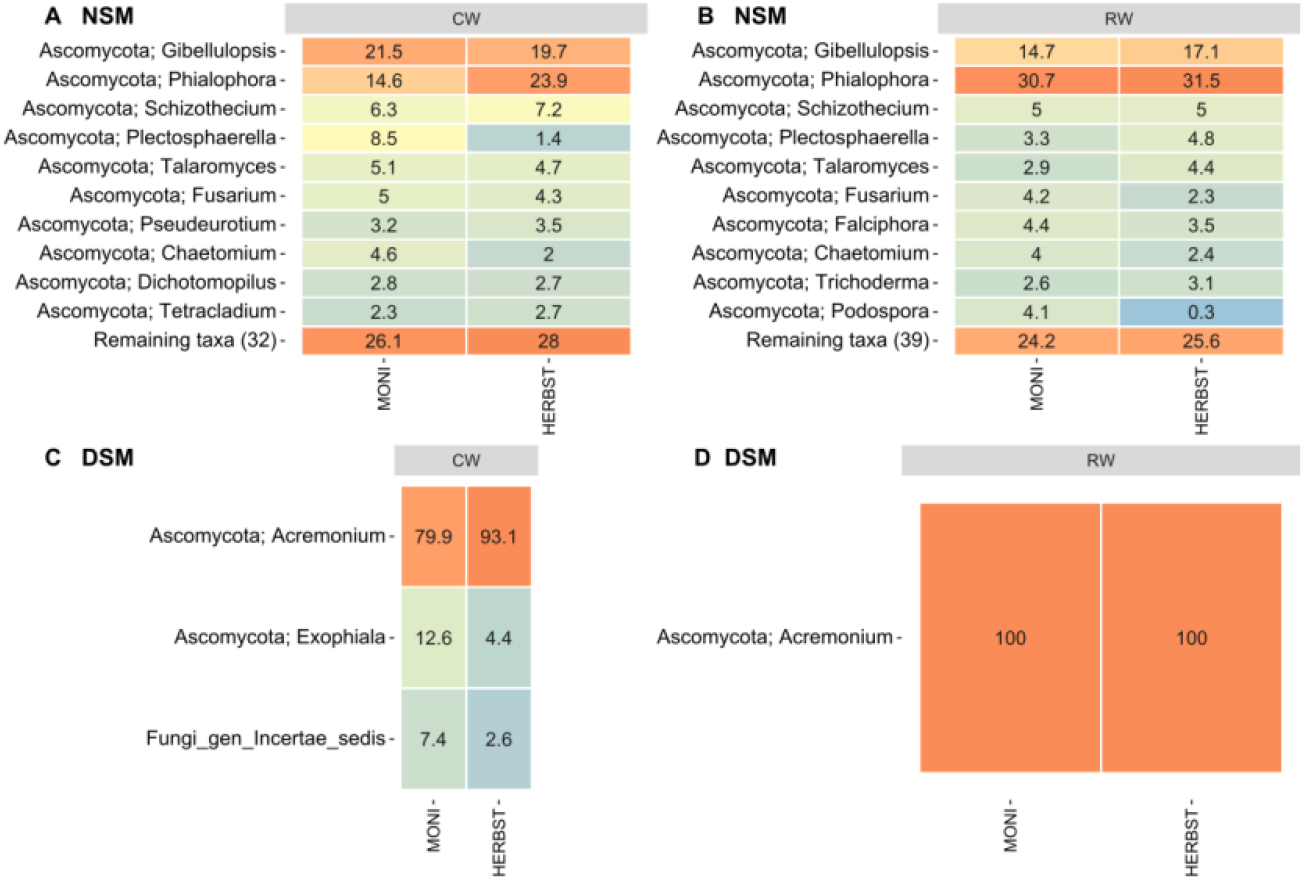
Fungal shared microbiome composition. Heatmap displaying the top 10 genera in the shared microbiome of cultivars MONI and HERBST grown in native soil with natural microbiome (NSM) and autoclaved soil with disturbed microbiome (DSM), under continuous (CW; A, C) and reduced (RW; B, D) watering. Numbers in the heatmap indicate the relative abundance of each taxon across sample collections.

## 4. Discussion

We investigated the rhizosphere microbiome of potato under reduced watering. To understand the influence of plants on the response of the rhizosphere microbiome, we used two cultivars of potato with different drought tolerance. Furthermore, we manipulated the soil reservoir microbial biomass, diversity and composition by soil autoclaving.

Interestingly, soil pre-treatment had no significant influence on growth parameters of both cultivars under continuous watering. However, the combination of DSM treatment with reduced watering significantly reduced the root length in the drought sensitive cultivar HERBST but did not affect root length of the drought resistant cultivar MONI. Roots play a critical role in plant adaptation to abiotic stress including water acquisition, nutrient uptake to the plant and interaction with microorganisms in the rhizosphere (Khan et al., 2016; Zinta et al., 2022). Furthermore, increased productivity of plants under drought has been correlated with key morphological traits influencing the root length and surface area of root systems (Comas et al., 2013). An inverse relationship (i.e., the larger the root system, the smaller the yield decline) has been observed under drought, suggesting that cultivars with deep root systems may be drought resistant (Zarzyńska et al., 2017). Previously, Boguszewska-Mańkowska et al. (2020) showed that GWIAZDA and TAJFUN, two drought tolerant cultivars of potato developed elongated roots when challenged with drought, whereas in drought susceptible cultivars (OBERON and CEKIN), the root length was unchanged under drought. These observations are not consistent with our results, potentially due to the choice of cultivars, the magnitude of the drought stress, pot size and the plant growth stage at the time of the treatment (tuber initiation). Potato responses towards drought have been already reported to vary with phenological stage, genotype, and stress severity (Gervais et al., 2021; Monneveux et al., 2013). In line with our results from the cultivar HERBST, a reduction in root length under drought has been shown for potato in another study (Albiski et al., 2012) However, this reduction was only visible when plants were cultivated in the soil with strongly reduced microbial diversity, suggesting a potential positive correlation between soil microbiome diversity and root growth.

### 4.1 Response of the rhizosphere microbiome to water reduction is cultivar dependent

Many studies have previously demonstrated that soil acting as the microbial reservoir is the main contributor to the rhizosphere microbiome (Inceoǧlu et al., 2012; Veach et al., 2019; Yang et al., 2022). This has been confirmed in our study by the strong impact soil pre-treatment had on the rhizosphere microbial communities (bacteria and fungi) of potato. Differences observed between NSM and DSM rhizosphere microbiomes can be explained by their respective source soils. In fact, soil autoclaving reduced the starting microbial biomass (C_mic_ and N_mic_) (**Supplementary Table S2**), microbial diversity (**Supplementary Figure S1**), and induced changes in the microbial composition (**Supplementary Figure S2**). In addition, soil autoclaving increased DOC and inorganic N (NH4+) (**Supplementary Table S2**), which may drive a complete restructuring of microbial communities as discussed in another chapter below. Nevertheless, it is important to emphasize that this study did not intend to investigate the effect of soil autoclaving on the subsequently acquired rhizosphere microbiome. We rather used this technique to manipulate the diversity of the natural soil microbiome in order to study the interaction between two potato cultivars of different drought tolerance and differently diverse communities of rhizosphere microbiota under reduced soil moisture.

#### Soil with natural microbiome (NSM)

Increasing evidence supports the role of root associated microbial communities in plant responses to environmental stresses. We hypothesized that when drought tolerance is mediated in potato by the recruitment of beneficial microbes, the rhizosphere microbiome would be affected in a cultivar-specific manner under reduced soil moisture, with the resistant cultivar MONI exhibiting a more adaptive microbiome under drought than the drought susceptible cultivar HERBST (**H1**). In the native soil with natural microbiome, MONI showed a stable rhizosphere microbiome regardless of the water regimes and potentially recruited less but drought tolerant microbes such as Rhizobiales (Santos-Medellín et al., 2017) under reduced watering. Unexpectedly, we witnessed a significant response to the reduced watering in HERBST rhizosphere microbiome (diversity and composition), resulting in potential recruitment of well-known drought-responsive taxa mainly in the Actinobacteriota phylum, and the root endophytic fungus *Falciphora* (Sun et al., 2020). Thus, although there is a host-dependent differentiation in the rhizosphere microbiome under reduced watering, we found no support for **H1**. Previously, Gaete et al., (2021) thoroughly selected tomato cultivars for drought tolerance and compared their rhizosphere microbiome under full and deficit irrigation regimes. As observed for MONI, the tolerant tomato cultivar showed unchanged bacterial and fungal diversity irrespective of the irrigation treatment, which the authors attributed to a buffering effect exerted by the tolerant cultivar on its rhizosphere (Gaete et al., 2021). In the same study, a significant increase in diversity during deficit irrigation was shown for the sensitive tomato cultivar, consistent with our observations in the HERBST rhizosphere. Specialized metabolites produced by *Solanum* plants, especially α-tomatine in tomato and α-solanine in potato (Friedman, 2006), two glycoalkaloids with known antimicrobial effects (Friedman, 2002; Milner et al., 2011) have been suggested to reduce the growth of many bacterial families in soil (Nakayasu et al., 2022). Increased microbial α-diversity suggests that reduced watering could interfere with the production of these metabolites in drought sensitive cultivars of Solanaceae.

Cultivar-dependent response to reduced watering observed in this study may be attributed to root exudates, considered the main factor in shaping the rhizosphere microbiome (Bulgarelli et al., 2013; de Vries et al., 2020; Dennis et al., 2010; Shi et al., 2011). Root exudates do not only depend on plant species and genotypes (Gschwendtner et al., 2011; Micallef et al., 2009), their quantity and quality may also change under drought conditions (Canarini et al., 2016; Naylor and Coleman-Derr, 2018; Preece and Peñuelas, 2016). Organic acids in the root exudates were found to increase under moderate drought conditions (Song et al., 2012) which, in turn stimulated microbial activities in the rhizosphere (Macias-Benitez et al., 2020). Another study established a positive correlation between exuded salicylic acid, GABA and Actinobacteria (Badri et al., 2013). However, the active recruitment of monoderm bacteria, such as Actinobacteriota under drought as shown elsewhere (Naylor et al., 2017; Santos-Medellín et al., 2017, 2021; Xu et al., 2018) and confirmed in HERBST rhizosphere needs to be further studied. Our results on potato cultivars of different drought tolerance give further support that tolerant or sensitive plants may engage with different microbial players when exposed to the same stress, suggesting a link between host-microbial interaction and the plant capacity to respond to drought (Gaete et al., 2021).

Regarding microbial taxa that potentially responded to the water regimes, two ASVs belonging to the order Rhizobiales were detected in MONI cultivated under reduced watering. Enrichment of Rhizobiales under drought was previously shown for rice (Santos-Medellín et al., 2017). Rhizobiales have been found as drought-responsive taxa in sugarcane rhizosphere (Liu et al., 2021) and furthermore, promotion of root development under drought has been attributed to these bacteria (Liu et al., 2021). Actinobacterial enrichment under drought has been observed in several studies across different plant species and environments (Naylor et al., 2017; Santos-Medellín et al., 2017, 2021; Xu et al., 2018). The Actinobacteria genus *Streptomyces* was found to be differentially abundant in the rhizosphere of drought-stressed potato (Faist et al., 2023). This is in line with our observation in the rhizosphere of HERBST, but not MONI. Xu et al. (2018) showed that drought increased colonization of *Streptomyces* isolates in the rhizosphere and roots of *Sorghum bicolor*, which according to the authors promoted plant root development during the stress. Plants protect themselves from oxidative stress through release of reactive oxygen species (Huang et al., 2017), whose damage can be effectively mitigated in plants by many species of *Streptomyces* (Lee et al., 2005; Leirós et al., 2014) to promote root growth for example (Voothuluru and Sharp, 2012; Xu et al., 2018). Besides some *Streptomyces* species being plant pathogens (Kopecky et al., 2019; Wei et al., 2022), our results from HERBST indicate that the recruitment of this genus in the plant rhizosphere may potentially have positive implications for plant fitness especially root growth under reduced soil moisture. In addition to *Streptomyces*, other drought enriched Actinobacteriota such as *Glycomyces* (Naylor et al., 2017; Yang et al., 2020)*, Marmoricola* (Gebauer et al., 2022), *Aeromicrobium* (Ma et al., 2022), *Mycobacterium* (Boukhatem et al., 2022) were found as potential responders to the reduced watering in the rhizosphere of HERBST. These taxa could act in synergy with *Streptomyces* or through different mechanisms to promote root growth. Under various stress, plants select in their rhizospheres beneficial microorganisms (Naylor and Coleman-Derr, 2018). Their mode of actions is generally attributed to the modulation of phytohormone levels (Afgan et al., 2016; Lugtenberg and Kamilova, 2009; Vacheron et al., 2013) such as ethylene which plays a critical role in plant responses to stress, especially at the root level (Mattoo and Suttle, 2017; Tanimoto et al., 1995). Many drought-responsive Actinobacteria contain the *acdS* gene (Gebauer et al., 2022) encoding the enzyme 1-amino cyclopropane-1-carboxlate (ACC) deaminase, which efficiently degrades the direct stress ethylene precursor (ACC) to ammonia and α-ketoglurate, thereby reducing ethylene biosynthesis under drought. Amplicon-based analysis of *acdS* gene showed an enrichment of the Actinobacteria *Marmoricola* in the rhizosphere of drought stressed barley (Gebauer et al., 2022), suggesting that the reduction of ethylene biosynthesis may be associated with root growth in HERBST, as this mechanism has been demonstrated under drought, flooding, heat, cold, pathogen colonization etc. (Gamalero and Glick, 2012). Root growth stimulated by auxin producing microorganisms is another mechanism reported to alleviate drought stress in plants (Bhattacharyya et al., 2021). An isolate from the root endophytic genus *Falciphora* co-cultivated with *Arabidopsis thaliana* improved lateral root growth via regulation of Auxin biosynthesis, signalling and transport in plant (Sun et al., 2020). While we identified *Falciphora* in the HERBST rhizosphere under reduced watering we have no indication of a positive modulation of root growth. However, the presence of this genus underlines the putative role of root endophytes in plant growth particularly under abiotic stress (Malicka et al., 2022). Potato is often considered as a drought-sensitive crop, mainly attributed to its shallow and sparse root system (Yuan et al., 2003). So far, our results suggest that both cultivars grown in the native soil potentially recruited beneficial microbes that potentially support root growth under reduced soil moisture. We found support for reports on drought tolerance of fungal communities generally being high in that only few fungal taxa responded to the reduced watering (Bazany et al., 2022). However, those taxa that responded to the reduced soil moisture were obviously influenced by drought either directly (saprotrophs) or indirectly by their root associated lifestyle (Lozano et al., 2021) suggesting a differential drought tolerance also in fungal communities as reported by Buscardo et al., (2021).

#### Soil with disturbed microbiome (DSM)

We hypothesized that when the rhizosphere microbiome plays a dominant role in drought resistance of the two potato cultivars, then a reduction of the soil microbial diversity would result in lower drought tolerance under reduced watering in both cultivars (**H2**). Surprisingly, despite the reduction of microbial diversity in the source soil, the rhizosphere microbiome (bacteria and fungi) of the resistant cultivar MONI was unaffected (alpha and beta diversity) by the reduced watering regime. Furthermore, a restricted number of bacterial biomarkers of reduced watering was found in the rhizosphere of MONI, as previously observed in NSM soil. This stability in the rhizosphere could have supported growth parameters of MONI and perhaps increased its leaf dry weight. However, the drought sensitive cultivar HERBST showed significant changes in bacterial community under reduced watering when plants were grown in the soil with reduced microbial diversity. Moreover, the latter induced the recruitment of other potential microbial responders such as Proteobacteria, Acidobacteriota and fewer Actinobacteriota under reduced watering. Acidobacteriota generally decrease in relative abundance under drought (Naylor et al., 2017; Santos-Medellín et al., 2017), suggesting that they are drought sensitive phylum. We argue that this substitution, however incomplete, of drought tolerant Actinobacteriota by Proteobacteria and Acidobacteriota could be one of the reasons behind the significant reduction in root length of HERBST cultivated under reduced watering. Therefore, since the plants were differentially affected by reduced watering in the soil with reduced microbial diversity, we reject **H2**. Interestingly, we were able to find that in the two potato cultivars, none of plant growth parameters was affected under continuous watering, despite the significant effect of soil pre-treatment on rhizosphere microbial communities. This suggests that plants would interact more with their rhizosphere microbiome under reduced soil moisture than under sufficient water supply.

In contrast to the bacterial community, fungi were not affected by water regimes and no potential responsive taxa were detected in the rhizosphere for either cultivar in DSM soil. In line with our observations, previous studies reported a less pronounced and even non-existent effect of drought on the structure of fungal communities in soil and root associated microbiomes (Barnard et al., 2013; Furze et al., 2017; Naylor et al., 2017; Ochoa-Hueso et al., 2018). This may partly be attributed to fungal spores which are highly persistent towards drought stress. In this respect also Streptomycetes and other Actinobacteria must be discussed as it cannot be excluded that their high relative abundance in the drought affected soils is partly related to their potential to form inactive forms, which cannot be differentiated from vegetative cells using an DNA based metabarcoding approach. Fungal communities in our study were less diverse than bacterial communities in the native soil, as previously showed elsewhere (Bazany et al., 2022; Gaete et al., 2021), and that the soil pre-treatment had almost eradicated them from the plant rhizosphere. This could lead to the non-identification of water responsive fungal taxa in the two cultivars, which we suggest may increase the susceptibility of HERBST exposed to the reduced watering.

### 4.3 Cultivars had a shared microbiome across soil pre-treatment and water regimes

Our study reveals the existence of shared microbiomes across the different sample groups providing evidence of a common response given by the two cultivars to the different treatments. Regarding the bacterial community, it appears that Actinobacteriota are the main drivers of this common response to the two water regimes in the native soil. In addition to the reasons mentioned earlier in this discussion, these bacteria have physiological (degrading recalcitrant compounds, sporulation) and structural characteristics (thicker peptidoglycan cell wall) (Naylor and Coleman-Derr, 2018) that allow them to thrive in dry environments. Thus, their strong interaction with the host rhizosphere may participate in the alleviation of stress in the two potato cultivars. *Gaiella* (Siebielec et al., 2020), *Nocardioides* (Liotti et al., 2018), *Arthrobacter* (Banerjee et al., 2010), *Marmoricola* (Gebauer et al., 2022) and *Streptomyces* (Faist et al., 2023) were the most shared Actinobacteriota. *Gaiella* was found very abundantly in soil with reduced moisture and is involved in nitrogen cycling (Siebielec et al., 2020). The other genera have been reported as plant growth promoting rhizobacteria with ability to produce auxins, siderophore and many other compounds (Boukhatem et al., 2022). Members of the fungal genera *Falciphora* often associated with an endophytic lifestyle (Malicka et al., 2022) and *Trichoderma* with strain specific drought tolerance (Singh et al., 2022) increased in relative abundance and have been previously recognized to encompass drought tolerant strains that potentially could sustain plant growth under stressful conditions. However, this assumption needs further investigations.

We also found that compared to the drought sensitive HERBST, the rhizosphere of the cultivar MONI showed a relatively stable number of bacterial core ASVs irrespective to soil pre-treatment and water regimes. Another striking finding was that autoclaved soil with reduced microbial diversity led to a complete restructuring of the shared microbial communities of the rhizosphere, making Proteobacteria and other fast-growing bacteria the main drivers to the water regimes. In fact, a drastic decrease in the relative abundance of Actinobacteriota, Chloroflexi, Acidobacteriota among many others, and a strong increase in Proteobacteria, Bacteroidota, Verrucomicrobiota and Firmicutes were observed. This suggests that the cultivar MONI, having the ability to recruit Proteobacteria and Verrucomicrobiota members as shown by the LEfSe analysis in NSM soil, was able to adapt quickly in the DSM soil even though it was exposed to the reduced watering. HERBST, preferring Actinobacteriota, was not able to establish an interaction as strong as that observed in the NSM soil under reduced watering. Furthermore, drought-responsive fungal taxa found in NSM soil were completely lost in the shared microbiomes in DSM soil. Collectively, the observed reduction in Actinobacteriota combined with that of drought tolerant fungi in the shared microbiomes may explain reduced root length of the less performing HERBST under reduced soil moisture.

## 5. Conclusion

In summary, this work illustrates how the strong impact of initial diversity and community composition in the rhizosphere influenced the response of differently drought tolerant potato cultivars to reduced soil moisture. Therefore, our study contributes to improve reduce knowledge gaps on potato and its rhizosphere microbiome interactions under reduced soil moisture. Using reduced soil microbiomes may provide an approach to address mechanisms that in drought susceptible potato cultivars prevent beneficial or neutral plant-microbe interactions. In addition, it should be further explored to which extent natural microbiomes support potato plants under drought by including more potato cultivars, different soils, and more severe drought scenarios. Although our results on potato support suggestions to include soil-root-microbe interactions in breeding programs towards drought tolerance, a number of spore-forming taxa was found in the microbiome. This points out the need of using metagenomic and metatranscriptomic approaches in future studies to address the activity (or the active implications) of microbial communities in the rhizosphere of potato plants under drought.

## Author contributions

BM, VR and MS designed the experiment of the study. KT and DM prepared the plant material used for the study. BM conducted the experiment and the laboratory work. BM and RS wrote the pipeline script and conducted the data analysis. BM, VR, KP and MS wrote sections of the manuscript. All authors contributed to manuscript revision, read, and approved the submitted version.

## Acknowledgments

The authors kindly thank the technicians Gudrun Hufnagel and Cornelia Galonska of the Research Unit for Comparative Microbiome Analysis at Helmholtz Munich for their support in the laboratory work.

