## Supplementary material for "Low soil moisture induces recruitment of Actinobacteria in the rhizosphere of a drought-sensitive and Rhizobiales in a drought-tolerant potato cultivar": Supplementary document.html

**Supplementary Figures**

**Table S2:** Measurement of the soil
microbial biomass (C_mic_ and N_mic_),
dissolved organic carbon (DOC), total dissolved nitrogen (TDN), inorganic
nitrogen (NH4-N, NO3-N), soil texture (Sand, silt clay), pH, and mineral
contents (P2O5, K2O, Mg) in the non-planted NSM, DSM prior to the experiment.
Mean values are reported in the table and statistical significance was
calculated using student t-test (∗p < 0.05, ∗∗p < 0.01, and ∗∗∗p < 0.001 and
\*\*\*\*p < 0.0001, n = 3). 

   
**Figure S1.**
Comparison of α-diversity based on Shannon index and observed species at the
start of the experiment of A) bacterial and B) fungal communities. 
Boxplots display the medians, tops and bottoms of the boxes represent 75th and
25th quartiles, and whiskers outside this range; dots illustrate the individual
observations in each sample group. A non-parametric Wilcoxon sum-rank test (p
< 0.05, n = 5) was applied to calculate significant differences across
sample groups and numbers above the boxes indicate the corresponding p-values. 

**Figure S2.** Ordination plots. Principal coordinate analysis (PCoA) of beta diversity at
the start of the experiment A) bacterial and B) fungal communities based on
weighted UniFrac distance. Colored shapes represent
soil pre-treatment (natural soil microbiome (NSM), green turkish
triangles and disturbed soil microbiome (DSM), brown circles). Soil
pre-treatment was separated along the first axis.  

**Figure S3.**  Composition of bacteria shared in the microbiomes of two potato
cultivars (MONI, HERBST) in natural (NSM) and disturbed soil (DSM) under
control (CW; A, C) and reduced watering (RW; B, D). Heatmap displays the top 20
shared bacterial microbiome members aggregated at the genus level. Numbers in
the heatmap indicate the relative abundance of each taxon across in each sample
type. 

**Supplementary Methods**

**Table S1:** Standard protocols used for
soil physico-chemical properties measurement

|  |  |  |
| --- | --- | --- |
| **Measurements** | **Supplier** | **Reference** |
| pH (CaCl) | AGROLAB Agrarzentrum GmbH | VDLUFA I, A5.1.1: 2016 |
| P_2_O_5_ | AGROLAB Agrarzentrum GmbH | VDLUFA I, A6.2.1.1: 2012 |
| K_2_O | AGROLAB Agrarzentrum GmbH | VDLUFA I, A6.2.1.1: 2012 |
| Mg | AGROLAB Agrarzentrum GmbH | VDLUFA I, A6.2.4.1: 1991 |
| Clay (<0.002 mm) | AGROLAB Agrarzentrum GmbH | DIN ISO 11277: 2002-08 |
| Silt (0.002 – 0.063 mm) | AGROLAB Agrarzentrum GmbH | DIN ISO 11277: 2002-08 |
| Sand (0.063 – 2 mm) | AGROLAB Agrarzentrum GmbH | DIN ISO 11277: 2002-08 |
